# Amplicon and metagenomic data from fumarole-associated geothermal features of Hawai□i

**DOI:** 10.64898/2026.02.26.708227

**Authors:** Jimmy H. Saw, Maximilian D. Shlafstein, Christina Pavloudi, Natalia Monsalve, Rebecca D. Prescott, Patrick S. G. Chain, Alan W. Decho, Stuart P. Donachie

**Affiliations:** Department of Biological Sciences, The George Washington University, Washington, DC, USA; University of Mississippi, Oxford, MS, USA; Los Alamos National Laboratory, Los Alamos, NM, USA; University of South Carolina, Columbia, SC, USA; School of Life Sciences, University of Hawai‘i at Mānoa, Honolulu, HI, USA; European Marine Biological Resource Centre-European Research Infrastructure Consortium (EMBRC-ERIC), Paris, France

## Abstract

The Hawaiian islands are among the most geologically and volcanically active places on Earth. While the Hawaiian Archipelago is known for its animal and plant diversity, much less is known about microbial diversity in the area’s diverse habitats. In this study, we focused on steam vent associated biofilms found on the most volcanically active island of Hawai□i, also known as the Big Island. From 46 samples from various biofilms and associated features around fumaroles emitting water steam, we generated amplicon and metagenomic sequences. Amplicon data showed that Chloroflexota and Cyanobacteriota are the numerically dominant phyla in these biofilm communities. We constructed 363 non-redundant medium to high-quality metagenome-assembled genomes (MAGs) that are at least 70% complete and with less than 5% contamination. Ten MAGs belong in the domain Archaea, and 353 belong in the domain Bacteria. This dataset could provide valuable insights into microbial diversity and ecology around volcanic features in Hawai‘i and elsewhere.

## Background & Summary

The Hawaiian Archipelago is considered to be a biodiversity hotspot, and also one of the most volcanically active places on Earth^1^. However, studies have shown that biodiversity in Hawai□i has been declining quickly^2,3^. Volcanic features found on the island – such as calderas, lakes, and fissures – are features that can form rapidly but could also disappear rapidly in this very dynamic environment. Microbial communities continue to thrive in these extreme yet dynamic environments, and their unexpected diversity has only become apparent in recently^4–6^. As features around volcanoes are often transient and can disappear rapidly, it is important to understand how rapidly such features are colonized, which species establish, the potential succession of the community, and what functions they might perform before these features disappear again. These habitats also serve as analog sites for understanding potential for life around volcanic systems on other planets. It is also notable that some of these volcanic features in Hawai□i host rare lineages that are important in our understanding of the early evolution of oxygenic photosynthesis^7,8^. Characterizations of microbes and microbial communities around volcanic habitats in Hawai‘i conducted previously focused on volcanic soils^9^, caves^5^, lakes^10^, fumaroles^4,5,11,12^, and submarine habitats^10,13,14^. However, most of these studies either employed clone libraries and Sanger sequencing of 16S rRNA or did not perform whole-genome shotgun sequencing of the samples.

In this study, we focus on the biofilms found near fumaroles on the island of Hawai□i, specifically those that emit water vapor instead of other gases such as sulfur dioxide. While few studies have investigated microbial communities around these fumaroles, to our knowledge this is the first to employ higher-throughput sequencing methods (such as Illumina) combining both amplicon and shotgun metagenomic sequences to characterize microbial diversity in these biofilms ^4,6,11,12,15^. Here, we used Illumina short-read sequencing techniques to characterize microbial communities around steam vent biofilms to generate amplicon (16S and 18S rRNA) and whole-genome shotgun (WGS) sequences. WGS data is important for investigating the potential functional roles microbes play in these communities. During 2019, we collected samples from biofilms or soil found near rock walls, surfaces, fissures, crevices, or caves in the Kīlauea caldera and the East Rift Zone (ERZ) that were all subjected to steam venting from directly below^5^. Steam from these features are generated by hot rocks below ground, which vaporize downward-percolating rainwater. We did not measure the composition of the steam and cannot exclude the presence of other gases in these steam. Samples were collected from a cave known as Big Ell in Kīlauea Caldera (in Hawaii Volcanoes National Park, HVNP), and from steam vents in the East Rift Zone (ERZ) that are publicly accessible (**Figure 1**).

**Figure 1.**
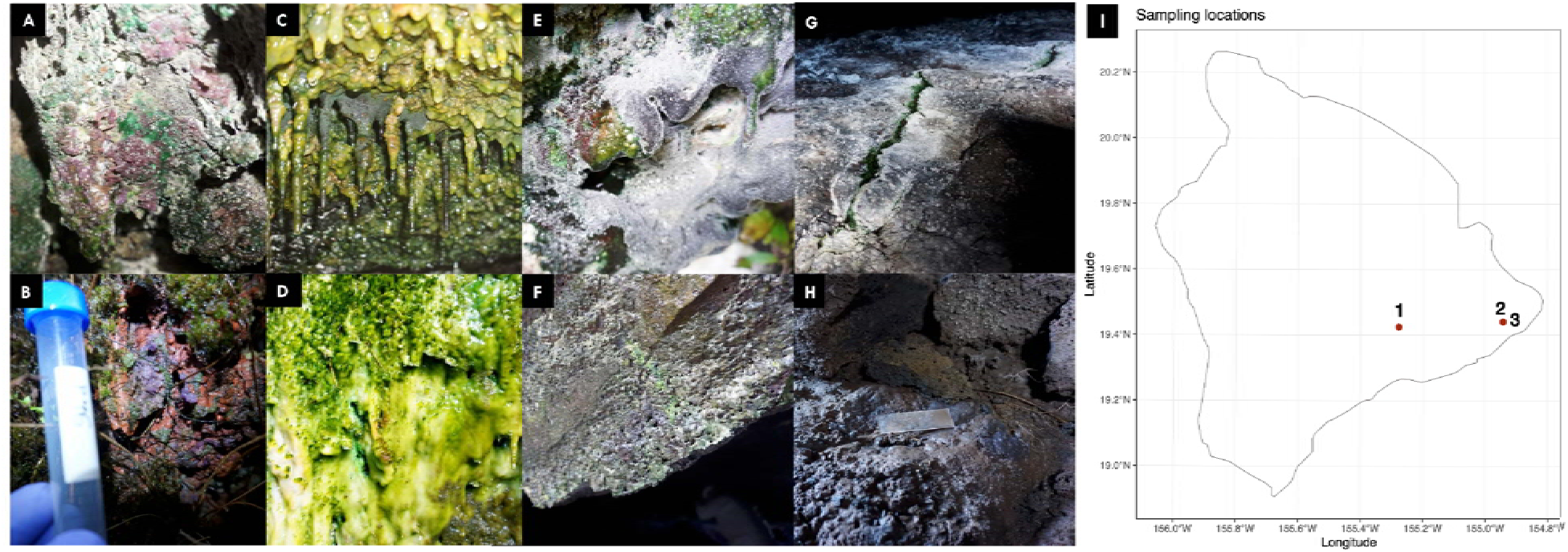
Images and a map showing sampling locations. (**A-B**) Purple biofilms dominated by *Gloeobacter* inside a pit-like feature inside the ERZ, (**C-E**) green to dark green/purple biofilms found in an open area within the ERZ with various steam vents surrounding it, (**F**) dried up biofilm inside a cave named “Big Ell” within Kīlauea Caldera inside Hawaii Volcanoes National Park, (**G**) dark green biofilms inside a fissure within Big Ell cave, (**H**) soil samples inside Big Ell, map showing sampling sites (1 – cave inside Kīlauea caldera, 2-3 – fumaroles within the ERZ).

Most of the samples were collected from epilithic biofilms on rock walls, crevices, or cave-like features near steam vents (**Figure 1** and **supplemental document**). Two distinct features in the ERZ showed notable characteristics: one of the two sites visited was a pit-like area where steam billowed from a main pit, and from smaller crevices in the vicinity of that pit. Notably, locals use it as a sauna and it is publicly accessible from Highway 130 after a short hike. Samples collected from that pit have the prefix “P” (except the sample that is named PDB5, which was collected from another site). Light intensity within this pit-like feature is less intense than other sites around it and most of the biofilm materials found in this “P” site showed purple coloration (**Figure 1 A-B** and **supplemental document**), reminiscent of the purple biofilm found near the entrance of Big Ell, a cave that is located in Kīlauea caldera where *Gloeobacter kilaueensis* JS1 was first isolated^7^. It is also notable that the purple-colored biofilms were mostly found in crevices or cave-like features that were sheltered from direct sunlight.

Biofilms found in another site in proximity to the “P” site were in a more open area exposed to higher light intensities and appeared mostly green (**Figure 1 C-E** and **supplemental document**). This site features an open area surrounded by multiple rock walls exposed to steam exiting the ground, roughly 100-150m away from the pit-like site. Samples collected from this area were assigned names starting with prefix “S”; one sample “PDB5” (i.e, mentioned above) also came from this site. It is important to note that there are numerous steam vent features in this area, but we were not able to collect samples from all of them due to limited accessibility of features of interest. Temperatures of the biofilms ranged from ∼30°C to 69°C (Supplemental Table **S1**).

Samples were also collected from Big Ell cave in the Kilauea Caldera. These included the dried remnants of epilithic biofilms we first sampled in 2009^16^, and designated here as “W” (**Figure 1F**). Dark green epilithic biofilms, labeled “V”, were collected from steam-emitting fissures (∼60°C) in the cave floor (**Figure 1G**), and those labeled “HW” were collected from soil-like material in the vicinity of where we deployed a sterile glass slide in 2009 (**Figure 1H**).

From the 46 biological samples collected, a total of 110 amplicon (16S and 18S rRNA) and WGS samples were generated (**Table 1** and Supplemental Table **S1**), representing a total of ∼1.8 billion raw sequencing reads (276 Gbp of raw sequences). WGS samples originating from related biofilms or features were co-assembled using metaSPAdes^17^, resulting in 20 metagenome assemblies. This generated a total of ∼16 million contigs representing 11 Gbp of assembled sequence data. Contigs larger than 1500 bp were then binned into 1307 metagenome-assembled genomes (MAGs) using Metabat2^18^ and further dereplicated using dRep^19^ to obtain 363 unique medium to high-quality MAGs that are at least 70% complete and with less than 5% contamination (**Figures 2 and 3**). According to the MIMAG guidelines^20^, 257 of the MAGs belong to the “high-quality” category, i.e., at least 90% complete and less than 5% contaminated.

**Table 1.** Sample metadata and accession numbers. N.D.: Not determined. *: 16S sequences generated using both universal and archaeal primers.

| Sample | Biosample Accession | Secondary Accession | Location | Temp (°C) | Data generated |
| --- | --- | --- | --- | --- | --- |
| HW02 | ERS27818420 | SAMEA120589944 | 19.424806 N 155.276578 W | N.D. | 16S*, WGS |
| HW12 | ERS27818421 | SAMEA120589945 | 19.424806 N 155.276578 W | N.D. | 16S*, WGS |
| HW13 | ERS27818422 | SAMEA120589946 | 19.424806 N 155.276578 W | N.D. | 16S*, WGS |
| P2 | ERS11952021 | SAMEA14358361 | 19.440835 N 154.942505 W | 38 | 16S, WGS |
| P3 | ERS11952022 | SAMEA14358362 | 19.440835 N 154.942505 W | 38 | 16S, 18S, WGS |
| P4 | ERS11952023 | SAMEA14358363 | 19.440835 N 154.942505 W | 38 | 16S, WGS |
| P8 | ERS11952024 | SAMEA14358364 | 19.440835 N 154.942505 W | 32 | 16S, WGS |
| P9 | ERS11952025 | SAMEA14358365 | 19.440835 N 154.942505 W | 32 | 16S, WGS |
| P12 | ERS11952026 | SAMEA14358366 | 19.440835 N 154.942505 W | 32 | 16S, WGS |
| P14 | ERS11952027 | SAMEA14358367 | 19.440835 N 154.942505 W | 31 | 16S, WGS |
| P15 | ERS11952028 | SAMEA14358368 | 19.440835 N 154.942505 W | 31 | 16S, WGS |
| P16 | ERS11952029 | SAMEA14358369 | 19.440835 N 154.942505 W | 31 | 16S, WGS |
| P18 | ERS11952030 | SAMEA14358370 | 19.440835 N 154.942505 W | 30 | 16S, WGS |
| P19 | ERS11952031 | SAMEA14358371 | 19.440835 N 154.942505 W | 30 | 16S, WGS |
| P20 | ERS11952032 | SAMEA14358372 | 19.440835 N 154.942505 W | 30 | 16S, WGS |
| P23 | ERS11952033 | SAMEA14358373 | 19.440835 N 154.942505 W | 38 | 16S, WGS |
| P24 | ERS11952034 | SAMEA14358374 | 19.440835 N 154.942505 W | 38 | 16S, WGS |
| P25 | ERS11952035 | SAMEA14358375 | 19.440835 N 154.942505 W | 38 | 16S, WGS |
| P27 | ERS11952036 | SAMEA14358376 | 19.440835 N 154.942505 W | 35 | 16S, WGS |
| P28 | ERS11952037 | SAMEA14358377 | 19.440835 N 154.942505 W | 35 | 16S, WGS |
| P29 | ERS11952038 | SAMEA14358378 | 19.440835 N 154.942505 W | 35 | 16S, WGS |
| S2 | ERS11952039 | SAMEA14358379 | 19.440786 N 154.941894 W | 43 | 16S, WGS |
| S3 | ERS11952040 | SAMEA14358380 | 19.440786 N 154.941894 W | 43 | 16S, WGS |
| S4 | ERS11952041 | SAMEA14358381 | 19.440786 N 154.941894 W | 43 | 16S, WGS |
| S7 | ERS11952042 | SAMEA14358382 | 19.440786 N 154.941894 W | 50 | 16S, WGS |
| S8 | ERS11952043 | SAMEA14358383 | 19.440786 N 154.941894 W | 50 | 16S, WGS |
| S13 | ERS11952044 | SAMEA14358384 | 19.440786 N 154.941894 W | 43 | 16S, WGS |
| S14 | ERS11952045 | SAMEA14358385 | 19.440786 N 154.941894 W | 43 | 16S, WGS |
| S15 | ERS11952046 | SAMEA14358386 | 19.440786 N 154.941894 W | 43 | 16S, WGS |
| S18 | ERS11952047 | SAMEA14358387 | 19.440786 N 154.941894 W | 55 | 16S, WGS |
| S20 | ERS11952048 | SAMEA14358388 | 19.440786 N 154.941894 W | 55 | 16S, 18S, WGS |
| S22 | ERS11952049 | SAMEA14358389 | 19.440786 N 154.941894 W | 47 | 16S, WGS |
| S23 | ERS11952050 | SAMEA14358390 | 19.440786 N 154.941894 W | 47 | 16S, WGS |
| S25 | ERS11952051 | SAMEA14358391 | 19.440786 N 154.941894 W | 51 | 16S, WGS |
| S29 | ERS11952052 | SAMEA14358392 | 19.440786 N 154.941894 W | 47 | 16S, WGS |
| S30 | ERS11952053 | SAMEA14358393 | 19.440786 N 154.941894 W | 47 | 16S, WGS |
| S31 | ERS11952054 | SAMEA14358394 | 19.440786 N 154.941894 W | 47 | 16S, WGS |
| S34 | ERS11952055 | SAMEA14358395 | 19.440786 N 154.941894 W | 43 | 16S, WGS |
| S35 | ERS11952056 | SAMEA14358396 | 19.440786 N 154.941894 W | 43 | 16S, 18S, WGS |
| S36 | ERS11952057 | SAMEA14358397 | 19.440786 N 154.941894 W | 43 | 16S, WGS |
| S37 | ERS11952058 | SAMEA14358398 | 19.440786 N 154.941894 W | 69 | 16S, WGS |
| PDB5 | ERS11952059 | SAMEA14358399 | 19.440786 N 154.941894 W | 43 | 16S, WGS |
| V2 | ERS11952060 | SAMEA14358400 | 19.424806 N 155.276578 W | 60 | 16S, 18S, WGS |
| W1 | ERS11952061 | SAMEA14358401 | 19.424806 N 155.276578 W | 30 | 16S, 18S, WGS |
| W3 | ERS11952062 | SAMEA14358402 | 19.424806 N 155.276578 W | 30 | 16S, WGS |
| W4 | ERS11952063 | SAMEA14358403 | 19.424806 N 155.276578 W | 30 | 16S, WGS |

**Figure 2.**
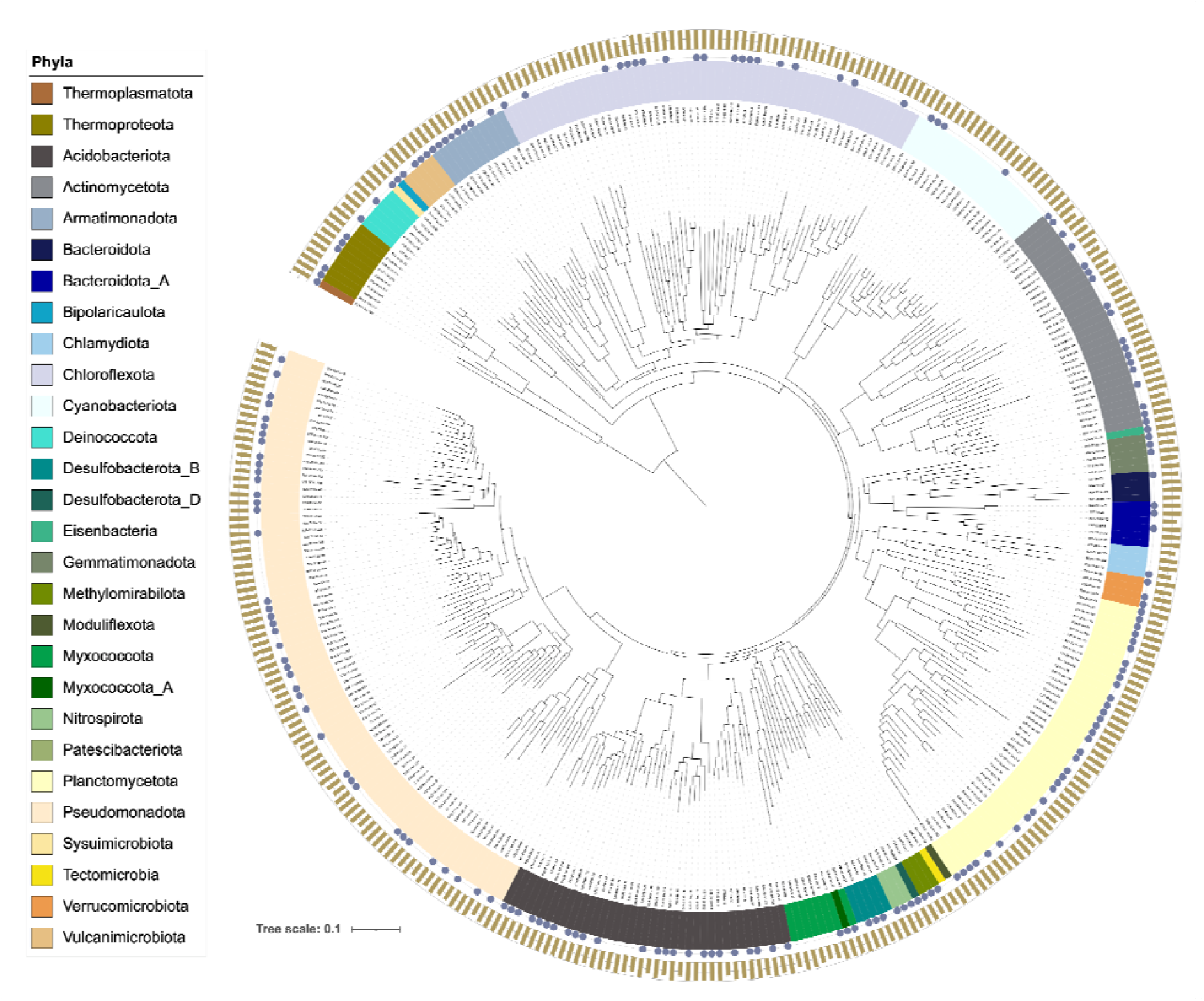
Maximum likelihood phylogenomic tree of 358 dereplicated archaeal and bacterial MAGs that are at least 70% complete and with less than 5% contamination. Phyla are colored and represented by the innermost ring next to the taxa labels. The presence or absence of circles in the middle ring indicate whether full or partial 16S rRNA gene fragments were recovered in the MAGs. The bar charts in the outermost ring show genome completeness estimates. A single MAG that belongs to the phylum Babelota was excluded by the PhyloPhlAn analysis pipeline and was not in this tree.

**Figure 3.**
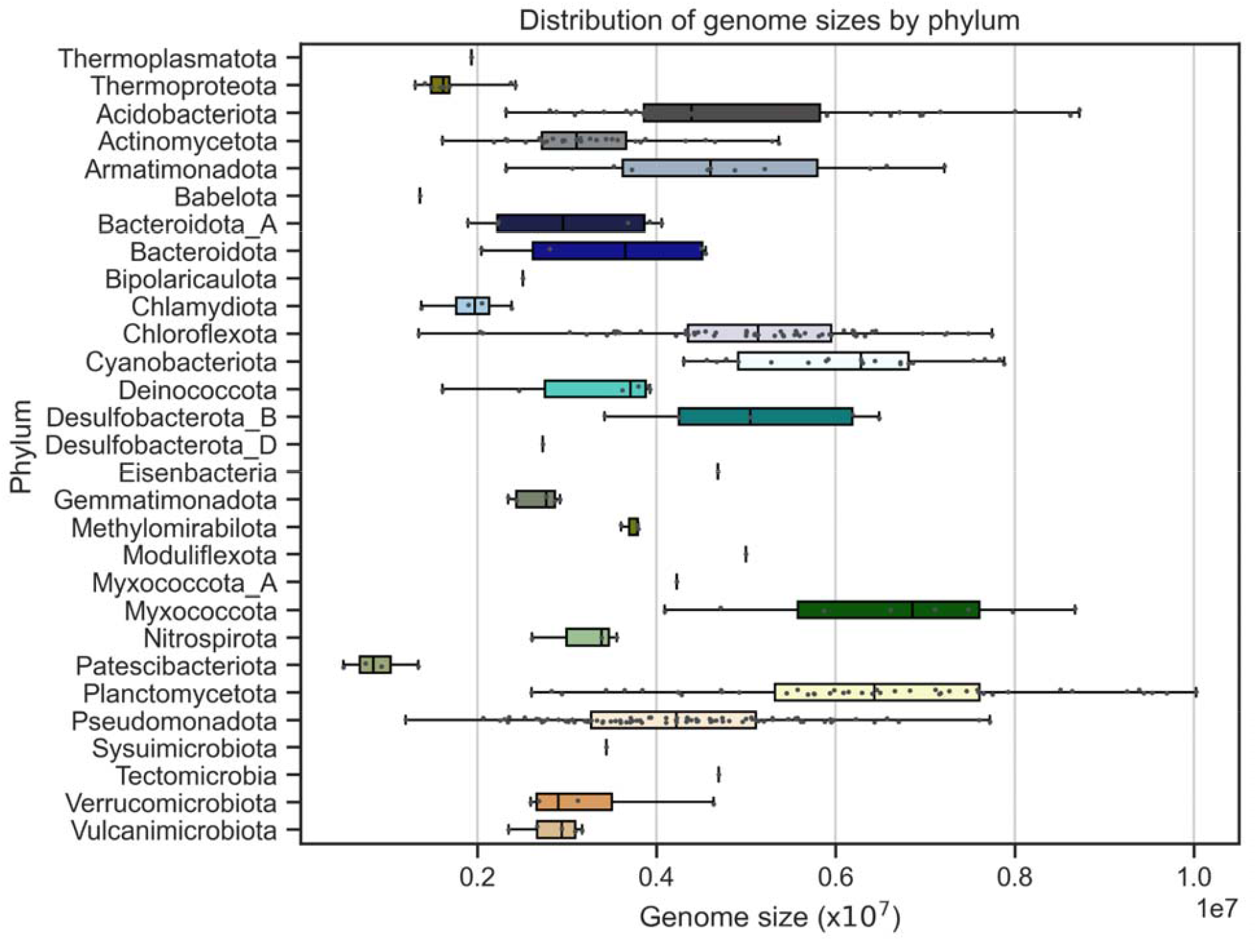
Boxplot showing distribution of genome sizes of the MAGs and whiskers showing minimum to maximum genome sizes of MAGs found in each phylum. Dots indicate each MAG in each phylum. Phyla are color coded using the same color categories used in Figure 2.

The MAGs were classified into 29 phyla using GTDB-Tk along with its reference database (release 10-RS226)^21,22^. Using PhyloPhlAn, we constructed a phylogenomic tree of all the dereplicated MAGs to show the total distribution of the MAGs by phylum, their completeness, and presence/absence of 16S sequences in the MAGs (**Figure 2**). Of the 363 unique MAGs, 10 belong to Archaea and 353 to Bacteria. Top 5 most numerous phyla represented among the MAGs are bacterial: Pseudomonadota (n=85), Chloroflexota (n=57), Planctomycetota (n=44), Acidobacteriota (n=39), and Actinomycetota (n=31). Largest genomes were observed in lineages that belong to Myxococcota, Planctomycetota, and Cyanobacteriota while the smallest genomes were observed in lineages belonging to Patescibacteriota, Thermoproteota, and Chlamydiota (**Figure 3**). Either full or partial 16S rRNA gene sequences were recovered from most of the MAGs we obtained (**Figure 2**). A single archeal MAG that belongs to the phylum Thermoplasmatota has a single contig of 1.93 Mbp with completeness score of 98%. We recovered three MAGs of non-photosynthetic cyanobacteria that belong to the class Vampirovibrionia among the MAGs. Their genome completenesses varied between 73.5 to 96.6% and genome sizes varied between 4.3-6.7 Mbp (Figures 2 and 3).

Most of the amplicon data were generated using universal primers targeting the V3-4 region of the 16S rRNA gene sequence but we also generated amplicons using Archaea or Eukarya-specific primers (against the 18S rRNA gene) for some of the samples (Supplemental Table **S2**). Amplicon data analysis using universal primers on rarefied samples showed that microbial communities of biofilms collected in proximity to fumaroles were highly diverse but numerically dominated by bacteria belonging to the Chloroflexota and Cyanobacteriota phyla (**Figure 4**). As expected, purple biofilm samples found in the pit-like feature had a high number of amplicon sequence variants (ASVs) that belong to the genus *Gloeobacter* (Supplemental data on Zenodo at DOI number: 10.5281/zenodo.18703655). Surprisingly, we detected a total of 1866 ASVs that belonged to the genus *Gloeobacter* despite having recovered only one unique MAG that was classified as *Gloeobacter kilaueensis* (Supplemental Table **S2**). The second most numerically dominant cyanobacterial order found in the biofilms (both P and S sites) is Cyanobacteriales with 2366 ASVs. We also detected a large number of Ktedonobacterales ASVs (n=5075) within the phylum Chloroflexota. By far, the most numerous ASVs were found within the order Rhizobiales (belonging to Pseudomonadota phylum) with a total of 7885 ASVs. Another notable order with a large number of ASVs in our dataset is Gemmatales (phylum Planctomycetota) with a total of 4871 ASVs. Tables displaying all ASVs found, their abundances in each sample, their classifications, and sequence fasta files have been deposited to Zenodo at DOI number: 10.5281/zenodo.18703655.

**Figure 4.**
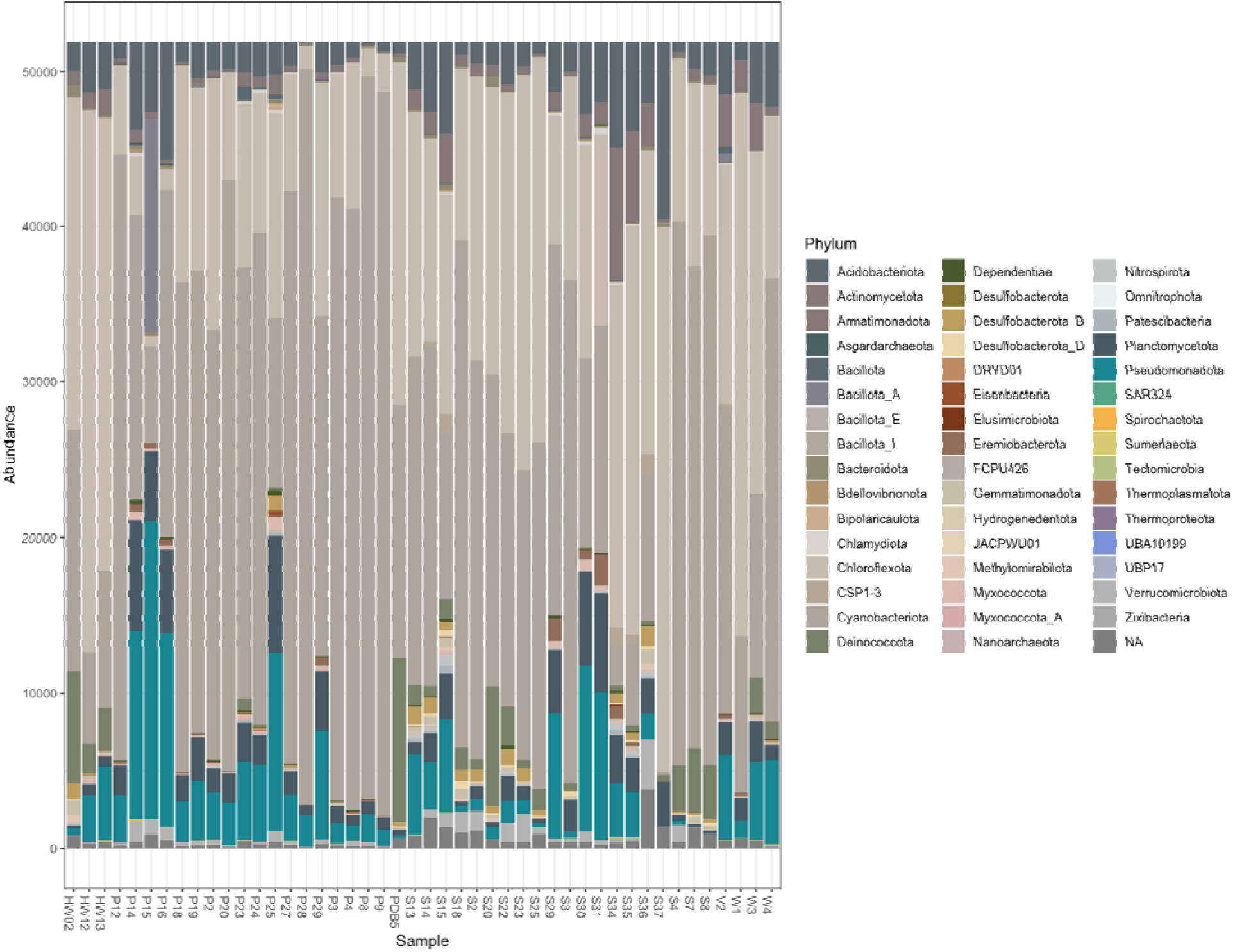
Stacked bar graphs showing archaeal and bacterial phyla identified and their abundances in each of the 46 samples. Samples were rarefied (sampled without replacement) to show an equal number of sequences in each sample (samples were ordered alphabetically). Taxonomic classification of the amplicon sequence variances (ASVs) was done using GTDB database release r220.

To the best of our knowledge, this dataset represents the largest dataset characterizing microbial communities occurring near features associated with steam vents or fumaroles in proximity to the volcanic regions of Hawai□i. Amplicon data provide a comprehensive survey of microbial diversity and abundance in these biofilms and the metagenomic data provides insights into functional roles these microbes play in these habitats. The metagenomic data could also be used to reveal viral diversity in these microbial communities and how they may be influencing the genomic evolution of Archaea and Bacteria. Lastly, this dataset may lend insight into understanding the early- and late-colonizers of newly-formed features of volcanic regions, not just in Hawai□i but also globally and could help us monitor if microbial diversity around these habitats is mostly stable or are experiencing declines in similar fashions as animal and plant species found on the islands.

## Methods

### Sample Collection

Biofilm and soil samples from fumarole-associated features were collected in August 2019 and sample sites have been described elsewhere^5^. Approximate sampling locations and images are provided (**Figure 1**, and supplemental data online). Briefly, biofilm samples were collected from steam vents at three locations (two along a highway between Pahoa and Kalapana, and a third in Big Ell cave in Kīlauea Caldera in HVNP) into sterile 15 or 50 mL polypropylene centrifuge tubes, kept on cold packs during transport to a lodging site, and frozen immediately upon arrival. Samples were shipped on dry ice to a laboratory at the George Washington University (GWU) and kept in a 80°C freezer until DNA extraction. Volcanic soil samples were also collected from the same cave in Kīlauea caldera.

### DNA extraction and sequencing

DNA was extracted from each sample using the ZymoBIOMICS DNA extraction kit (Zymo Research, Orange, CA, USA) and quantified using Qubit 4 fluorometer and high-sensitivity dsDNA assay kit (Thermo Fisher Scientific, Waltham, MA, USA). Samples passing the concentration threshold needed for sequencing were sent to three different sequencing facilities: a) Los Alamos National Laboratory, b) Advanced Studies in Genomics, Proteomics, and Bioinformatics (ASGPB) at the University of Hawai‘i at Mānoa, and c) Center for Quantitative Life Sciences at Oregon State University (Supplementary Table S1). For amplicon sequencing, we used three sets of primers targeting the 16S and 18S rRNA genes: 16S universal primers targeting V3-4 regions (341F - CCTACGGGNGGCWGCAG and 785R - GACTACHVGGGTATCTAATCC)^23^, 16S Archaea-specific primers targeting the V3 region (349F - GYGCASCAGKCGMGAAW and 806R - GGACTACVSGGGTATCTAAT)^24,25^, and 18S primers targeting the V4 region (Uni18SF - AGGGCAAKYCTGGTGCCAGC and Uni18SR - GRCGGTATCTRATCGYCTT)^26^. Amplicon samples were sequenced on an Illumina MiSeq instrument, and whole-genome shotgun metagenomic sequencing was conducted on HiSeq 3000 (Oregon State University) or NextSeq 500 (Los Alamos National Lab) instruments (Illumina, Inc, San Diego, CA, USA).

### 16S rRNA amplicon data analysis

Raw amplicon sequences were processed with cutadapt (version 2.9)^27^ to remove the primers. The reads were then analyzed with the DADA2 package (version 1.20)^28^ in R (version 3.5.2). Taxonomic assignment was performed with GTDB database release 220^22^ for the 16S rRNA data and SILVA database version 132 for the 18S rRNA data^29^. Samples were rarefied to the lowest common denominator (sampling without replacement) so that an equal number of reads were used to plot diversity and abundance bar charts. The Phyloseq (version 1.36)^30^ and ggplot2 (version 3.3.5)^31^ packages in R were used for the creation of barcharts. Detailed workflow for the analyses were documented in a Github repository (https://github.com/SawLabGW/HawaiiSteamVents).

### Metagenome assembly, binning, and quality assessment

Raw metagenomic sequences were processed using BBDuk version 38.87^32^ with input parameters: ktrim=r, minlen=50, mink=11, tbo=t, rcomp=f, k=21, ow=t, zl=4, qtrim=rl, trimq=20, to remove Illumina adapter sequences and low quality regions. Quality filtering excluded regions with a score less than 20 and reads under 50 bp in length. FastQC version 0.11.9 was then used to assess the quality of the trimmed sequences before downstream processing. Sequences from the same sample sites were combined before assembly to improve sequencing depth. MetaSPAdes version 3.15.4^17^ was used to assemble the combined trimmed sequences using default parameters but with k-mers of 21, 33, 55, and 77. Seqtk version 1.3^33^ was used to first create a list of the assembled contigs over 1kb, which were then mapped back to the trimmed reads with BBWrap version 38.87^32^ using default parameters. For combined samples, contigs were individually mapped to each of the trimmed reads before combination. These mapped reads were sorted and indexed using Samtools version 1.10^34^ with default parameters. The JGI summarize script that was packaged with BBmap tool was then used to determine the depths of contigs over 1,500 bp, allowing for binning with Metabat2 version 2.15^18^ using default parameters. MAGs produced by Metabat2 were further dereplicated using dRep tool version 3.6.2 with default parameters^19^.

### Taxonomic classification of MAGs and phylogenomics

The GTDB-Tk version 2.4.1 was used to infer the taxonomy of all MAGs using the GTDB reference database release 226^21,22^ using the “classify_wf” and the following parameters: -- skip_ani_screen --no_mash. Barrnap version 0.9^35^ was used to identify 16S ribosomal RNA genes in all of the bacterial MAGs with parameters: --kingdom bac, --reject 0.05. Taxonomic assignment of the 16S genes was performed using blastn version 2.10.1^36^ and the SILVA database version 138.1^29,37^ parameter -evalue 1e^-5^. Phylophlan version 3.1.68^38^ was used to obtain the concatenated alignment of single-copy marker genes found in dereplicated MAGs. IQ-TREE version 3.0.1 was used to construct the maximum likelihood phylogenomic tree of all the MAGs using the LG+F+G4 model of amino acid substitution^39^. The resulting phylogenomic tree was visualized using iTOL version 7.4.2^40^ and annotated with completeness data from CheckM and presence/absence of 16S rRNA gene information as obtained by barrnap.

### Data Records

All raw 16S, 18S, and shotgun metagenomic sequences have been deposited in European Nucleotide Archives (ENA)^41^ under the project accession number PRJEB52128. Accession numbers for all raw sequences are provided in Table 1 and also in Supplementary Table S1.

### Technical Validation

We used FastQC to assess quality of raw amplicon and metagenomic sequences, Quast^42^ to assess quality of metagenome assemblies, and CheckM (version 1.1.3)^43^ to assess completeness and contamination of MAGs. We also used dRep^19^ to dereplicated redundant MAGs to only keep unique high-quality MAGs for further analysis. Our threshold for high-quality MAGs were genome completeness of at least 70% or more and contamination of 5% or less.

## Data Availability

All of the raw reads for amplicon and metagenomic data are available at European Nucleotide Archive (ENA) under the project accession number of PRJEB52128 (available at http://www.ebi.ac.uk/ena/data/view/PRJEB52128). Dereplicated MAGs and sequences of ASVs are available on Zenodo at DOI number: 10.5281/zenodo.18703655.

## Code Availability

All workflows or scripts used to analyze the data were reposited to a Github repository: https://github.com/SawLabGW/HawaiiSteamVents.

## Author Contributions

The study was conceived by JS and SD. JHS collected samples, provided funding to sequence them, analyzed data, and wrote the paper. MDS performed metagenomic assembly, binning, and classification of the MAGs. CP extracted DNA from some of the samples and analyzed the amplicon samples. NM extracted DNA from most of the samples. RDP, PSGC, and AWD provided funding to sequence some of the samples. SD collected samples and coordinated the sampling with rangers from the Hawai‘i Volcanoes National Park. JS wrote the first draft of the manuscript, and all authors contributed to the final version of the manuscript.

## Competing Interests

The authors declare no competing interests.

## Acknowledgements

We would like to thank Liliana deSmither and Harry Schick who helped us locate Big Ell, a lava cave in Kīlauea Caldera. The permit number for sample collection in the Hawaii Volcanoes National Park is HAVO-2009-SCI-0029. We gratefully acknowledge the computing resources provided on the High-Performance Computing cluster operated by Research Technology Services at The George Washington University.

## Funding

JS was supported by the startup funds and University Facilitating Funds (UFF) from The George Washington University, and NSF CAREER Award (number 2442122). RP was supported by an NSF Postdoctoral Fellowship in Biology (award number 1711856) and a NASA Postdoctoral Fellowship. AD was supported by a NASA Exobiology grant (award number 80NSSC18K1064). PSGC was supported by the US Department of Energy (award number LANLF59T).

## Notes

https://www.ebi.ac.uk/ena/browser/view/PRJEB52128

### Competing Interest Statement

The authors have declared no competing interest.

